# Disentangling age-dependent DNA methylation: deterministic, stochastic, and nonlinear

**DOI:** 10.1101/2020.10.07.329987

**Authors:** O. Vershinina, M.G. Bacalini, A. Zaikin, C. Franceschi, M. Ivanchenko

## Abstract

DNA methylation variability arises due to concurrent genetic and environmental influences. Each of them is a mixture of regular and noisy sources, whose relative contribution has not been satisfactorily understood yet. We conduct a systematic assessment of the age-dependent methylation by the signal-to-noise ratio and identify a wealth of “deterministic” CpG probes (about 90%), whose methylation variability likely originates due to genetic and general environmental factors. The remaining 10% of “stochastic” CpG probes are arguably governed by the biological noise or incidental environmental factors. Investigating the mathematical functional relationship between methylation levels and variability, we find that in about 90% of the age-associated differentially methylated positions, the variability changes as the square of the methylation level, whereas in the most of the remaining cases the dependence is linear. Furthermore, we demonstrate that the methylation level itself in more than 15% cases varies nonlinearly with age (according to the power law), in contrast to the previously assumed linear changes. Our findings present ample evidence of the ubiquity of strong DNA methylation regulation, resulting in the individual age-dependent and nonlinear methylation trajectories, whose divergence explains the cross-sectional variability. It may also serve a basis for constructing novel nonlinear epigenetic clocks.

## Introduction

It is well known that DNA methylation changes with age^1,2^, and its variability predominantly increases, as manifested both in cross-sectional^3 – 5^ and longitudinal data^5,6^. Still, the explanation of the nature of methylation heterogeneity with age remains a largely open question so far. Earlier results on examining twin cohorts indicate that the changes in methylation variability during aging are the product of both genetic (heritable) effects *G* and adaptation to (common) environmental influence *E, G× E*^7,8^. The dynamical perspective offers a somewhat different angle on variability as an outcome of the interplay between deterministic and stochastic *D × S* influences. The relative importance of these factors has not yet been determined. In the current study, we focus on the age-related variably methylated probes and attempt to discriminate whether the variability changes mainly deterministically or stochastically, and if deterministically, whether linearly or not.

We address the three main questions: (i) what is the relative contribution of deterministic and stochastic evolution to variability of DNA methylation with age? (ii) what is the relation between the age-dependent change in methylation level and its variance, if any? (iii) what are the mathematical functional types of age-dependence? Deterministic change of variability could originate from intrinsic and extrinsic heterogeneity. In the *G × E* context, genetic differences and environmental factors, such as climate, level of pollution, social and economical factors, determine individual aging trajectories. At the same time, *G × E* fluctuations like genetic or replication noise, random environmental cues might contribute to stochastic-like changes in DNA methylation. The resulting general age trend of methylation level can thus be accompanied by a completely different behavior of individual trajectories. Deterministic divergence can result from heterogeneity of initial conditions and parameters of the methylation change (e.g. slope and intercept) for each individual (cf. Figure 1-a). The high level of fluctuations can lead to a biased random walk behavior (see illustration in Figure 1-b). Longitudinal analysis is required to disentangle *D × S* processes.

**Figure 1.**
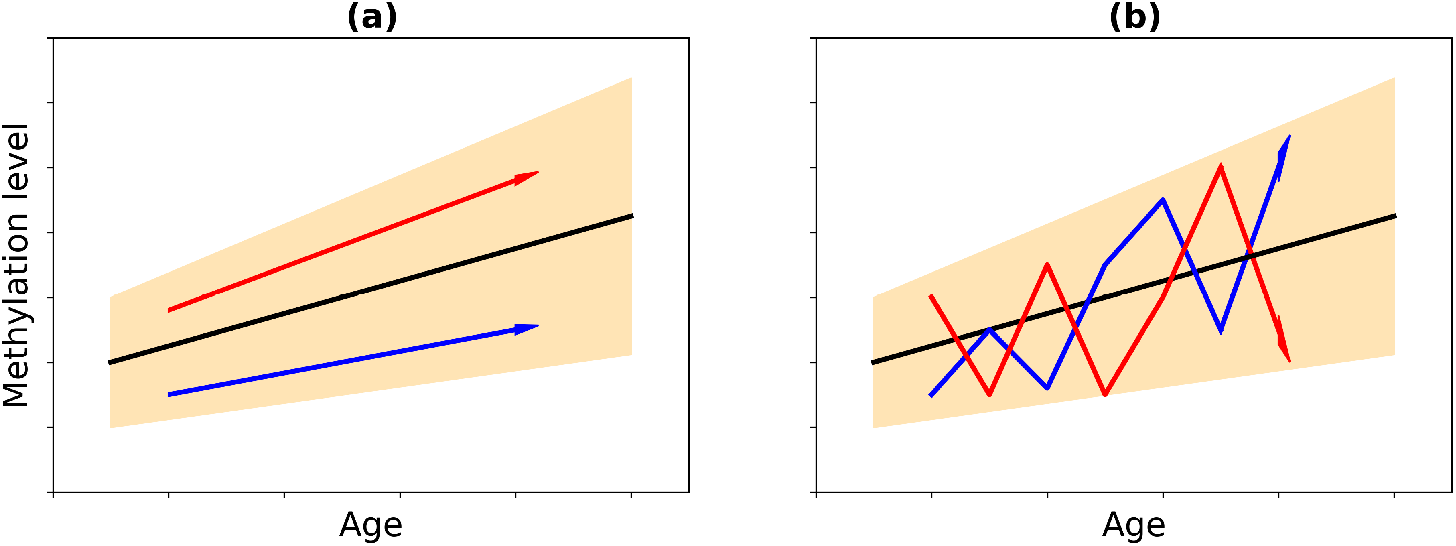
Illustration of possible mechanisms behind increasing variability with age: **(a)** deterministic divergence due to heterogeneity between individuals and **(b)** due to stochasticity.

Another important aspect is the determination of the law of methylation changes with age. Earlier studies were mostly aimed at identifying linear biomarkers of aging, that is, those CpGs whose methylation level varies linearly with age^9 –11^. Although it has been noticed that there are CpGs whose methylation changes logarithmically before adulthood^11^, CpG probes, whose methylation profiles differ from linear ones throughout aging were not found. In this regard, we conduct a systematic study and develop a procedure for finding nonlinear epigenetic biomarkers of aging. These are the novel biomarkers that have not been identified previously, since existing approaches are aimed at selecting the best linear predictors of age.

## Materials and Methods

### Datasets

We consider two DNA methylation datasets obtained using the Illumina Infinium HumanMethylation450 BeadChip on blood DNA: the cross-sectional methylation dataset of the Swedish population with identifier GSE87571^12^ from the Gene Expression Omnibus (GEO) repository^13^, and the longitudinal data from The Swedish Adoption/Twin Study of Aging (SATSA)^14^ from ArrayExpress repository^15^. The choice is motivated by their having the largest age span among the currently available cross-sectional and longitudinal datasets, that is critical for the study.

Raw data files were extracted and pre-processed using *minfi* Bioconductor package^16^. Samples having a detection p-value > 0.01 (indicating a poor quality signal) in more than 5% of probes were removed from the analysis, leaving 729 samples in GSE87571 dataset (341 males and 388 females aged 14 to 94 years) and 442 samples in SATSA dataset (179 males and 263 females characterized by 1 to 5 data points with an overall age span from 48 to 98 years). The SATSA dataset includes 180 pairs of twins (104 female-female, 74 male-male and 2 male-female). Raw data were normalized using the *preprocessFunnorm* function for GSE87571 and *preprocessQuantile* function for SATSA. Probes with a detection p-value > 0.01 in more than 1% of samples, probes mapping on sex chromosomes, probes with internal SNPs and cross-reactive probes^17^ were excluded from each dataset, leaving 414950 and 380137 probes in GSE87571 and SATSA datasets, respectively. Since previous studies indicate differential methylation between the two sexes^18–21^, we analysed males and females separately.

As known both beta and M-values are used in the literature for DNA methylation analysis, depending on the purpose of study. We set our choice on beta values to address its (non)linear age-dependence, divergence and also with regard to implications for epigenetic clocks^22,23^.

Additionally, blood cell counts were estimated from methylation data using Horvath’s online calculator^24^, which implements the method developed by Houseman et al.^25^ The residuals of methylation values were calculated from regression model for the dependence of methylation levels (beta values) on proportions of CD8T cells, CD4T cells, NK cells, B cells and Granulocytes separately for males and females. To eliminate negative values, residuals were additionally shifted up by a constant equal to the absolute value of the smallest residual. To avoid false effects due to the presence of different proportions of cells, the analysis performed for the beta values was completely repeated for the residuals.

### Estimation of deterministic and stochastic components of variability

First, we identify the probes, in which DNA methylation variability significantly changes with age (age-associated variably methylated positions, aVMPs) both in GSE87571 and SATSA datasets. For each dataset, we calculate the population-average variability of CpG sites as 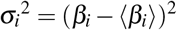, where ⟨*β*_*i*_ ⟩ is the average methylation level taken over a sliding window of 10 years, 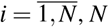 is the number of sites. Then we perform a statistical test on age dependence of 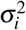, setting the null-hypothesis rejection p-value threshold to 0.001 for GSE87571 and 0.05 for SATSA after Benjamini-Hochberg correction^26^ (the difference in p-value thresholds accounts for the smaller age span of SATSA dataset, as compared to GSE87571).

Second, we disentangle *D × S* contribution to the variability growth with age by estimating the signal-to-noise ratio, *SNR*, for the selected aVMP probes based on the longitudinal dataset SATSA for both beta values and residuals. Further, to avoid repetition, all methods are described for beta values. The signal-to-noise ratio is the quantifier widely used in the field of signal processing and communication to assess the intensity of fluctuations in the time series data^27^. Specifically, it measures the ratio of average methylation, ⟨*β*⟩, to deviation, *σ*. For each probe we calculate three different types of *SNR*: for individual longitudinal trajectories; for twin pairs and for merged data points (“clouds”), when the longitudinal information is disregarded. For an individual trajectory *z* that has more than two methylation measurements at different age, we build the linear regression model, *β*_*r*_. Then we calculate deviation for each point according to *σ*_*j*_ = |*β*_*j*_ *−β*_*r, j*_ |, where 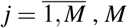 being the number of data points per person. The signal-to-noise ratio is defined as the ratio between the regressed methylation level to deviation at specific points, averaged along the trajectory, 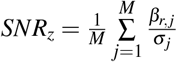. Finally, we obtain a single *SNR*^*trajectories*^ value for a CpG probe by averaging *SNR*_*z*_ over individuals.

To calculate the next type of *SNR*, we consider the methylation trajectories of twin pairs, for which the total observation interval contains at least two points. If one twin has a methylation measurement at some point in time, while the other does not, then the second is approximated by two neighboring points. We consider only pairs of twins of the same sex (two male-female pairs were excluded). For each pair of twins *t p* the following values are calculated: an averaged methylation 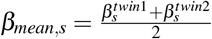 and a deviation 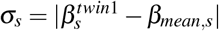, where 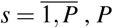 is the number of data points for a pair of twins.

In the last formula, it is not important which twin’s methylation is considered, since the average value *β*_*mean*_ at each time point is exactly in the middle between the methylation of twins. For each twin pairs we calculate the average signal-to-noise ratio as 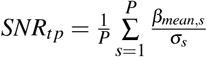 and obtain the final *SNR*^*twin−pairs*^ for each CpG site, averaging over all the pairs under consideration.

Alternatively, we pool methylation values together ignoring longitudinal and individual attributes, and calculate *SNR* for resulting sets of beta values. The average methylation value ⟨*β*⟩ is also taken over a sliding window of 10 years, the deviation is calculated as *σ*_*l*_ = |*β*_*l*_ *−*⟨*β* ⟩_*l*_ | for each subject and time point, 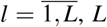 is the number of methylation points, and the signal-to-noise ratio for each CpG site is then 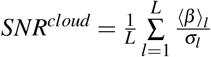. The ratio between the two quantities, 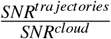, is the key quantifier, indicating the contribution of heterogeneity between individuals to the age-related increase of methylation variability. It is expected to be much bigger that 1 in the case Figure 1-a, and order 1 in the case Figure 1-b.

### Classifying age-related patterns of methylation level and variability

The dependence of CpG methylation levels *β* on age is fitted with the linear regression model in GSE87571 dataset. We restrict our attention to the CpG probes, which show significant age-associated methylation changes (age-associated differentially methylated positions, aDMPs), satisfying the criteria: 1) the linear regression slope is greater than 0.001, in other words, the probes, whose absolute methylation change on the span of 100 years would make at least |Δ*β* |= 0.1 (|Δ*β* | = 0.001 a year); 2) linear regression determination coefficient *R*^2^ is greater than the 99% percentile for the distribution of *R*^2^.

To identify CpG probes in which variability also significantly changes with age (age-associated differentially and variably methylated positions, aDaVMPs), we take the above selected aDMPs in male and female subsets. We calculate variability of each CpG site 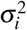, build the linear regression model for the dependence of 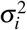 on age and set the null-hypothesis rejection value threshold to 0.001 (after Benjamini-Hochberg correction).

The list of aDaVMPs can be split into four classes, according to increasing/decreasing methylation level/its variability with age. Moreover, we investigate the potential functional dependence between methylation level and its variability and analyze the following characteristics^28^:

- Squared coefficient of variation^29^, 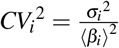 or 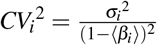, where 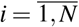
- Fano factor^30^, 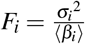 or 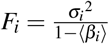, where 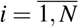.

Normalization to 1*—* ⟨ *β*_*i*_ ⟩is performed when the methylation level and variability show opposite trends with age. Then we perform statistical tests for the age dependence of 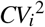 and *F*_*i*_, and identify the cases, when the null-hypothesis cannot be rejected at p-value threshold 0.001 (after Benjamini-Hochberg correction). If one or another normalization of variability removes age dependence, we infer a corresponding functional dependence of variability on the methylation level as *σ*^2^ *∼ β*^2^ or *σ*^2^ *∼ β*.

### Identification of probes having nonlinear age-associated methylation changes

Identification and discrimination of nonlinear vs linear trends requires the largest age span possible, so that we again focus on GSE87571 dataset. For the previously found list of aDMPs, we build linear regression models for linear and logarithmic beta values, *β*, and age: 1) *β* = *k · age* + *b*; 2) ln *β* = *α ·* ln(*age*) + *p*; 3) ln *β* = *γ · age* + *q*, where *k, b, α, p, γ, q* are fitting constants. Linear regression in the logarithmic axes corresponds to the power law fit 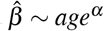, while regression in the semi-logarithmic axes corresponds to the exponential fit 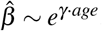. The quality of fits is estimated by coefficients of determination. For nonlinear fits the coefficients of determination *R*^2^ are calculated as 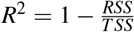, where 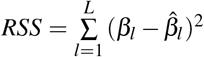 is the residual sum of squares, 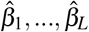 are fitted beta values, 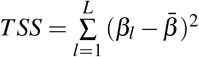 is the total sum of squares (proportional to the variance of the data), 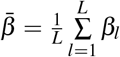 is the mean of the observed data, *L* is the number of subjects. In addition, we make regression for the complementary beta value, 1 *− β*, on age. If for a CpG site the determination coefficient of power law/exponential fit for the complementary beta value is at least 5% higher we take forth the former as a better model fit.

Ultimately, we compare the three coefficients *R*_*linear*_^2^, *R*_*pow*_^2^, *R*_*exp*_^2^. If one of the nonlinear fits has a determination coefficient at least 5% higher than that of the linear fit, then the age dependence of the probe is identified as nonlinear. Further on, if one of nonlinear fits has determination coefficient higher than the other, we identify the nonlinear dependence as the power law or exponential, accordingly.

For the remaining linear CpG sites, we also apply a “weak” criterion for nonlinearity. That is, if the determination coefficient of nonlinear fits (*R*_*pow*_^2^ or *R*_*exp*_^2^) lies within *±*5% of the determination coefficient of linear fit, *R*_*linear*_^2^. Such probes, for which both linear and nonlinear fits are equally suitable, make the majority of aDMPs.

The *methylgometh* function implemented in the *methylGSA* R package^31^ was used to calculate the enrichment of gene ontology terms in the selected lists of aVMPs, deterministic, stochastic and nonlinear probes. GO terms with an adjusted p-value < 0.01 were retained as significant.

## Results

### Identification of probes having age-associated methylation variability changes

To identify CpG probes that display age-associated methylation variability changes (age-associated variably methylated positions, aVMPs) we considered GSE87571 and SATSA Infinium 450k datasets, analyzing males and females separately. The choice of aVMPs is based on the construction of linear regression models for the dependence of the methylation variability, *σ*^2^, on age. First of all, we tested the age-dependence variability in GSE87571 dataset with the null-hypothesis rejection p-value 0.001. The resulting number of selected aVMPs was 37379 for males and 30420 for females. Among them we selected a subset that manifests age-dependent variability according to longitudinal SATSA dataset (at p-value 0.05, due to smaller age span of the dataset). In result, we obtained 9627 aVMPs for males and 2110 aVMPs for females (Supplementary File 1). Gene ontology (GO) analysis showed that both males and females aVMPs enrich ontologies related to development and ion transport (Supplementary File 2), as previously reported^21^. Interestingly, the number of aVMPs is noticeably greater for males, suggesting that methylation variability could be sex-specific.

We applied Fisher exact test to explore whether these aVMPs are enriched in specific genomic regions. It has turned out that the selected aVMPs probes show significant enrichment of CpG islands, north and south shores (p-values 6.48e-175, 1.79e-92, 6.1e-41 respectively) for males, while for females enrichment is restricted to north and south shores only (p-values 1.98e-68, 1.81e-38), see Figure 2-a. As for the part of the genome, aVMPs probes are significantly enriched in the 1stExon and TSS1500 (p-value 1.18e-05, 2.62e-24) for males and TSS1500 (p-value 5.76e-14) for females, see Figure 2-b. Also these sites are significantly enriched in the enhancer region with p-values 2.45e-08, 1.89e-08 for males and females, respectively.

**Figure 2.**
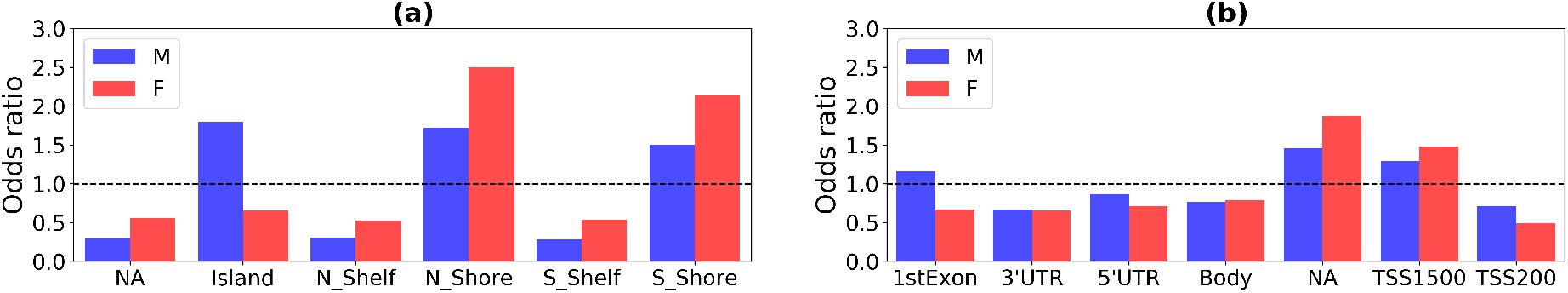
Enrichment (odds ratio) of genomic localizations for aVMPs calculated for males (blue) and females (red): **(a)** enrichment of genomic regions, **(b)** enrichment of genome parts.

We obtain qualitatively the same results for the residuals of methylation values obtained by regressing out the dependence of beta values on blood cells counts. In terms of numbers, 5692 aVMPs for males and 2077 aVMPs for females were identified (Supplementary File 3). These probes are significantly enriched in CpG islands, north and south shores (p-values 1.14e-03, 3.96e-88, 5.36e-46), TSS1500 (p-value 1.33e-26), enhancer region (p-value 1.42e-14) for males and for north and south shores (p-values 3.1e-58, 1.7e-41), TSS1500 (p-value 5.69e-09), enhancer region (p-value 7.16e-14) for females.

We compared the lists of CpGs obtained for beta values and residuals. The number of overlapping probes is 4100 for males and 1356 for females. Intersecting probes are marked with ‘X’ in the corresponding Supplementary Files 1, 3.

### Estimation of deterministic and stochastic components of variability

To investigate the deterministic and stochastic contribution to methylation variability, we focused on the longitudinal dataset SATSA. Primarily, for the previously identified aVMPs we calculated the average signal-to-noise ratio, *SNR*, for single (individual) trajectories. Larger *SNR* values are associated with the greater deterministic component in the age-related individual variability of methylation. The probability density functions (PDF) for the logarithms of *SNR* values are shown in Figure 3. To take into account the possible deterministic trend in the average methylation (cf. Figure 1), we also calculated the average *SNR* for the “clouds” of data points, that is *β* -values for all individuals and time points for a specific CpG, merged in a single set. The ratio between the two kinds of *SNR* characterizes the weight of deterministic component in individual longitudinal dynamics. Figure 4 shows the Manhattan plot for the logarithm of this ratio, 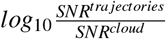.

**Figure 3.**
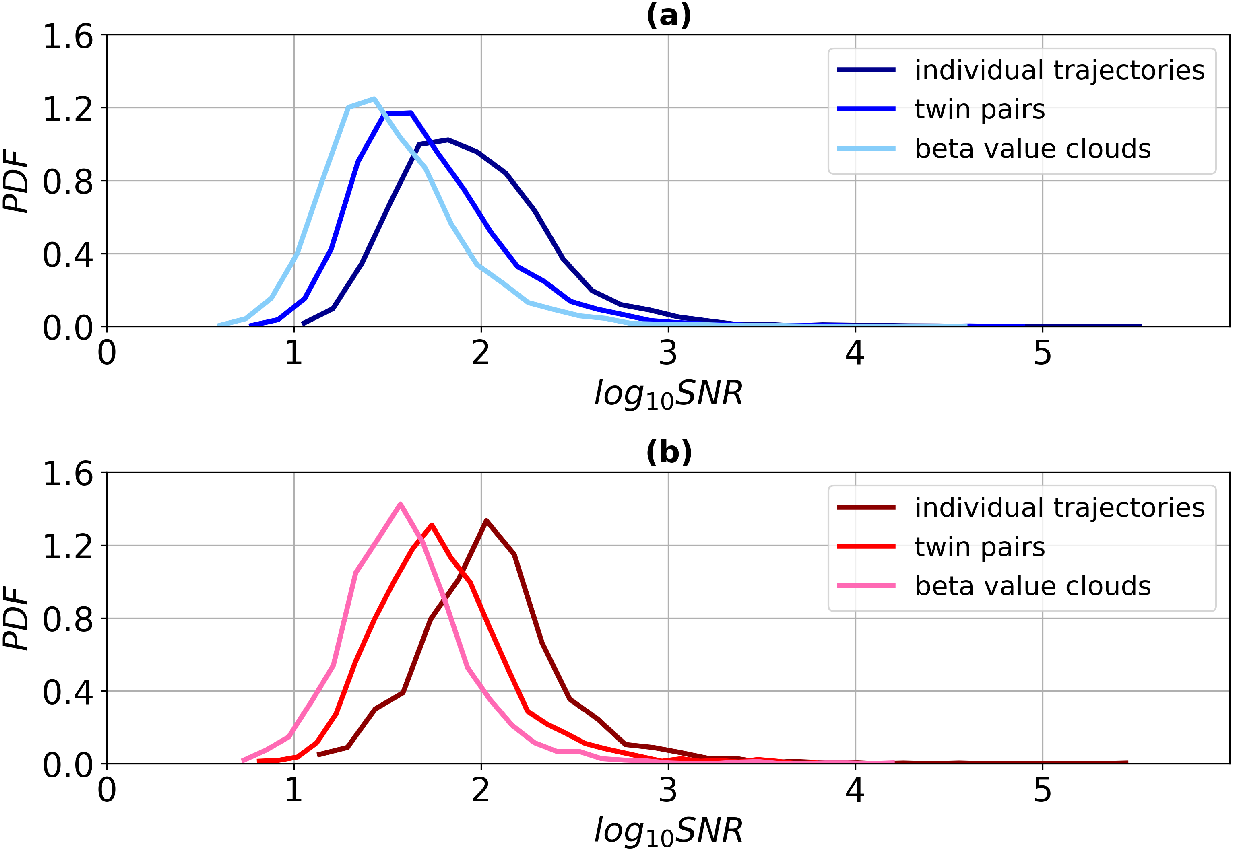
The probability density function (PDF) for *log*_10_ *SNR* of aVMPs. Top panel **(a)** - for males (dark blue - individual trajectories, blue - twin pairs, light blue - clouds of beta values), bottom panel **(b)** - for females (dark red - individual trajectories, red - twin pairs, pink - clouds of beta values).

**Figure 4.**
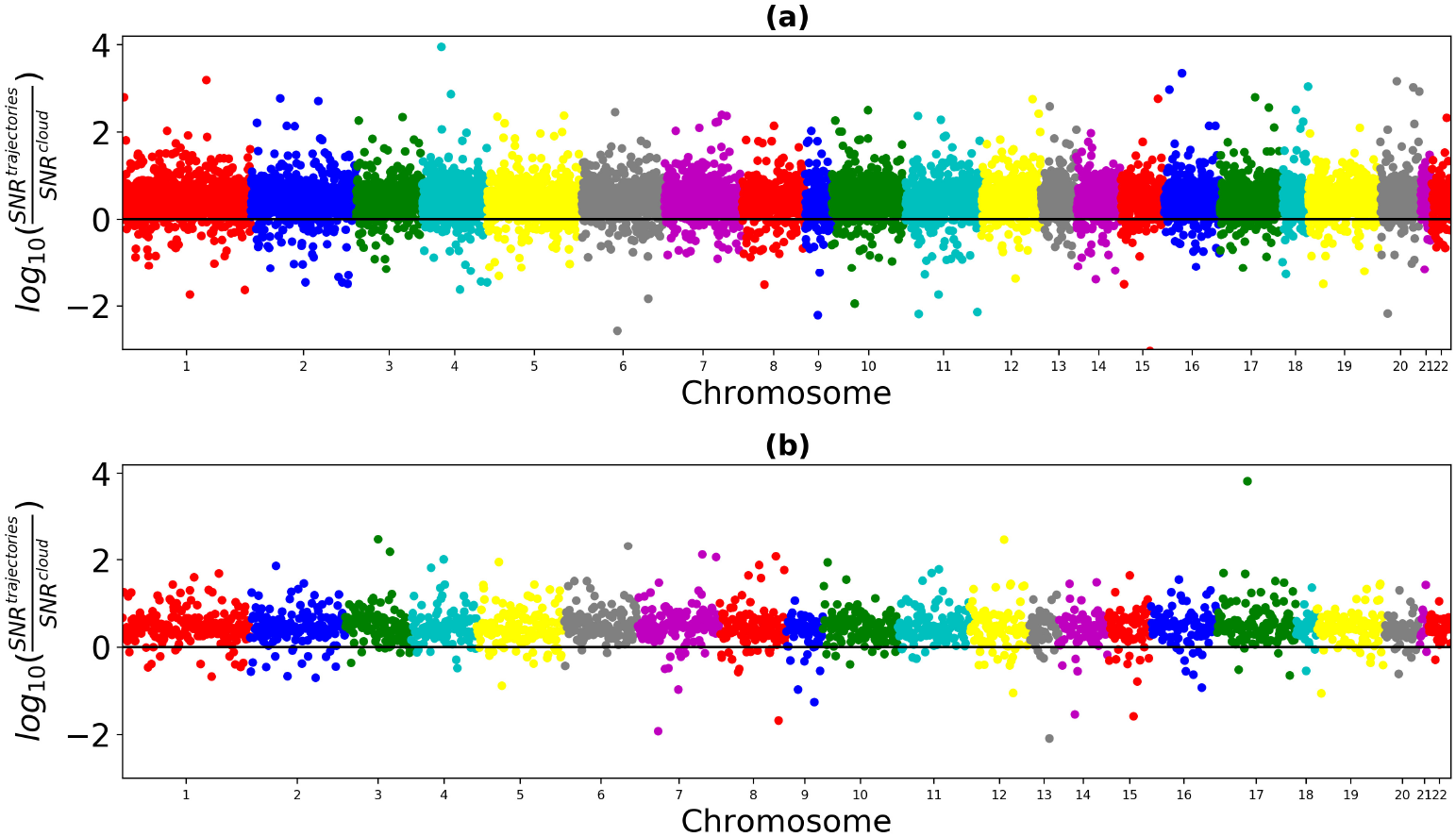
Manhattan plot of genome-wide logarithm of the ratio of the *SNR* of individual trajectories to the *SNR* of a cloud of beta values: top panel **(a)** – for males, bottom panel **(b)** – for females. The black lines are the zero value of the logarithm, that is, the situation when the *SNR* of individual trajectories and the *SNR* of a cloud of beta values are equal.

It follows, that for most of CpGs the value is positive (see Figure 4), and also PDF for individual trajectories remains to the right with respect to PDF of “clouds” (see Figure 3). That is, in most of the cases the average signal-to-noise ratio is greater for individual trajectories than for merged beta values. Therefore, methylation variability for individual longitudinal data changes substantially less with age, than the variability in the whole population. It suggests that the latter is dominated by population heterogeneity, individual differences at the origin and systemic environmental cues, which supports the major role of deterministic mechanism of age-dependent variability, sketched in Figure 1-a. The number of such CpGs amounts to 8722 CpGs (90.6%) for males and 1944 CpGs (92%) for females. The alternative picture is manifested for 905 CpGs for males and 166 CpGs for females, where the signal-to-noise ratio is greater for clouds of beta values. There, the stochastic mechanism of age-dependent variability appears dominant. aVMPs divided into deterministic and stochastic are presented in the Supplementary File 4 for males and females. GO analysis did not highlight ontologies specifically enriched by the subset of stochastic aVMPs: the 17 significantly enriched GO in males were all enriched also by deterministic probes, while the analysis of stochastic females aVMPs did not return significant results (Supplementary File 5 and Supplementary Figure 1).

Markedly, one observes sex differences in the *D × S* balance again, males exhibiting a stronger stochastic component in age-dependent methylation variability (see Figure 3), with the Kolmogorov–Smirnov test yielding p-value = 0.9925 for the PDF of *SNR* based on individual trajectories and p-value = 0.7753 when based on “clouds” of beta values.

To further illustrate our findings, we calculated the *SNR* for pairs of twins, which have a common genetic background and stayed in a common environment at least during childhood. Quite expectedly, the signal-to-noise ratio in the twins turned out to be higher than that of the beta value clouds (the curve of PDF lies to the right in Figure 3), and less than that of the individual trajectories. In other words, the relative variability between trajectories of twins is greater than the variability of a single individual trajectory, but less than the cross-sectional variability within the population.

Qualitatively the same results were observed when the analysis was performed on the residuals. In particular, the number of deterministic CpGs is 91.4% for males and 92.3% for females of the number of selected aVMPs. Similar lists of deterministic and stochastic probes for residuals are presented in Supplementary File 6. Common deterministic CpGs for beta values and residuals are marked with ‘X’, and common stochastic CpGs are marked with ‘XX’ in the corresponding Supplementary Files 4, 6.

### Classification of age-related methylation and variability changes

The pronounced deterministic component in the age-related changes of methylation variability raises the further question of studying the possible connection between the variability and average methylation level itself. To make the analysis conclusive, it is necessary to employ datasets with the age span as large as possible. In practice, one has to resort to cross-sectional datasets, among which GSE87571 emerged to be the only suitable choice at present.

In this subsection, we investigate the mathematical functional dependence between CpG methylation average and variability, as they change with age. Attention is focused on CpG sites that show significant age-associated methylation changes (age-associated differentially methylated positions, aDMPs, cf. Materials and Methods for details). The resulting number of selected aDMPs is 3827 for males and 3850 for females (Supplementary File 7). Further, we selected aDMPs with significant age-dependent methylation variability (age-associated differentially and variably methylated positions, aDaVMPs) and obtained 2075 probes for males and 2282 for females (Supplementary File 8). Using the similar analysis on the residuals we defined 2347 aDMPs for males, 2619 aDMPs for females (Supplementary File 9) and 592 aDaVMPs for males and 1008 aDaVMPs for females (Supplementary File 10). Probes that are common to beta values and residuals are marked with ‘X’ in the respective Supplementary Files 7-10.

We identified four classes of aDaVMP age dependence: 1) methylation and variability increase (1713 CpGs for males, 2133 CpGs for females); 2) methylation and variability decrease (24 CpGs for males, 13 CpGs for females); 3) methylation decreases and variability increases (330 CpGs for males, 133 CpGs females); 4) methylation increases and variability decreases (8 CpGs for males, 3 CpGs for females). The percentage of CpG probes in these classes is given in Figure 5, and typical examples are presented in Figure 6. More than 50% of aDaVMPs showed same direction changes in the level of methylation and variability during aging. Overall, apart from few exceptions, we observe an increase in variability with age, which is consistent with the earlier studies^3 – 5^. We find that decreasing variability is related to saturation towards complete (de)methylation (0 or 1 methylation levels).

**Figure 5.**
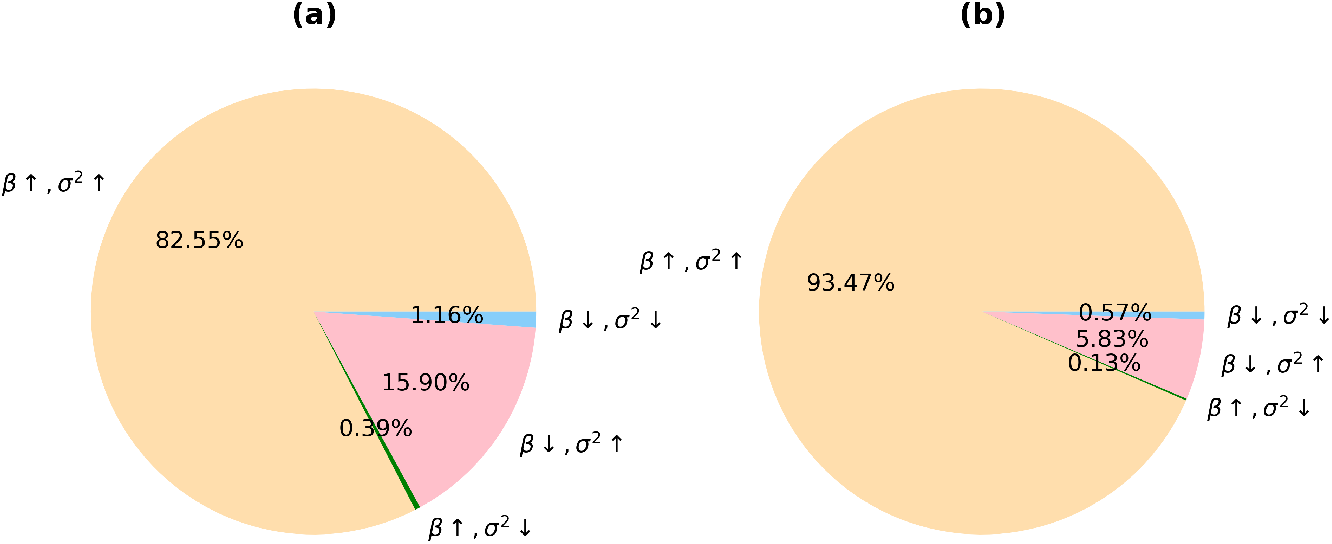
The percentage of aDaVMPs with different quality types of methylation and variability changes in GSE87571 dataset for males **(a)** and females **(b)**: case of methylation and variability increasing (beige); case of methylation and variability decreasing (lightblue); case of methylation decreasing and variability increasing (pink); case of methylation increasing and variability decreasing (green).

**Figure 6.**
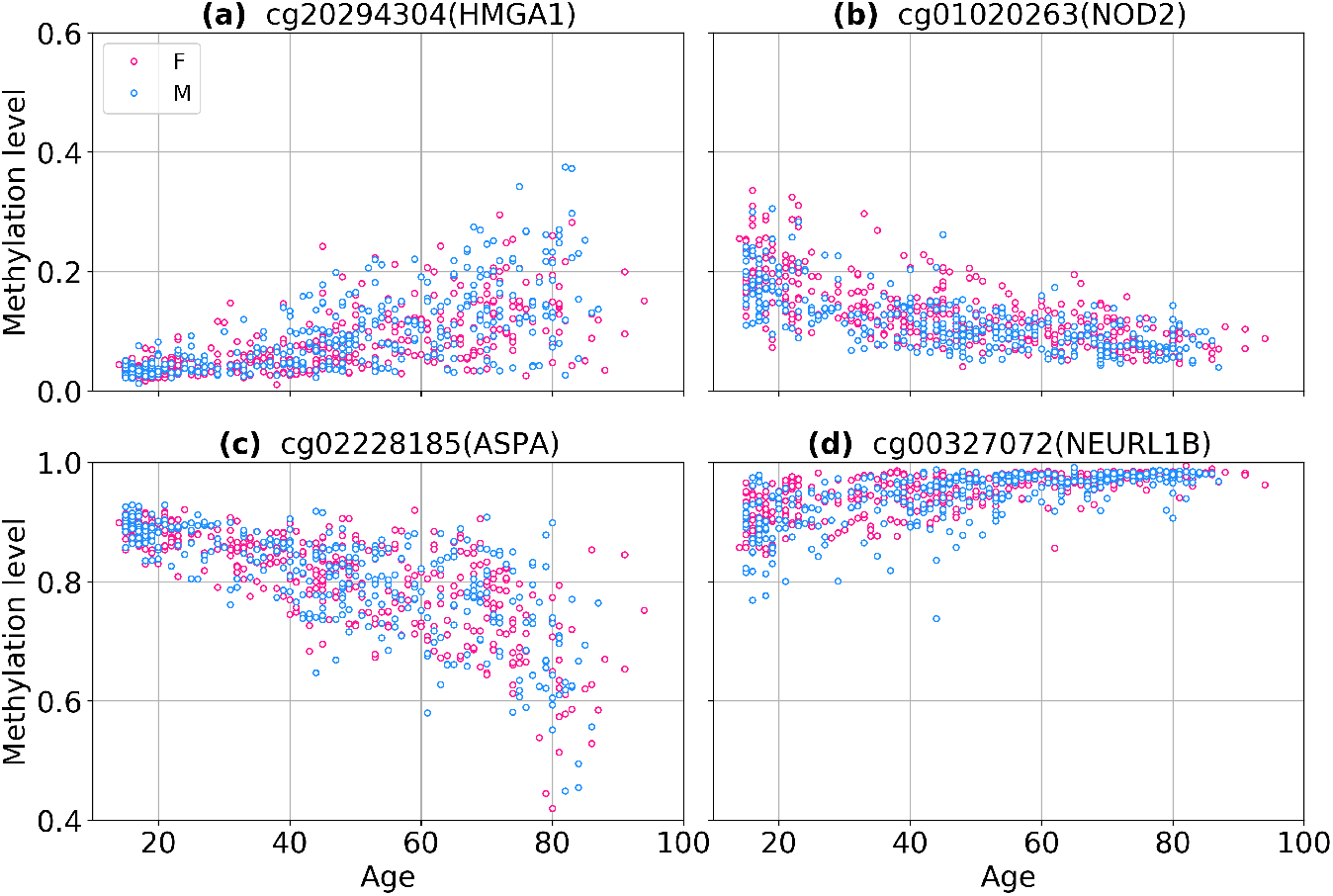
Examples of CpG probes having different quality types of methylation and variability changes in GSE87571 dataset: methylation and variability increase **(a)**, methylation and variability decrease **(b)**, methylation decreases and variability increases **(c)**, methylation increases and variability decreases **(d)**.

The studies of biological noise have highlighted several model functional dependencies between expectation of measurable and its variability, *σ*^2^ = *σ*^2^ (*β*)^28^. In order to probe for these model cases, we calculated the squared coefficient of variation^29^, *CV*^2^ = *σ*^2^ */β*^2^, and the Fano factor^30^, *F* = *σ*^2^ */β*, for each aDaVMP. It is worth noting that the former is independent on the number of trials (that can be associated with the time arrow), for example, for the biased random walk, and the latter for the Poisson process. We identified the probes when one or another normalization of variability removes age-dependence of *σ*^2^, and thus inferred its type (cf. Materials and Methods for details). That is, if *CV*^2^ for a CpG site does not depend on age, then *σ*^2^ *∼ β*^2^, and if *F* does not depend on age, then *σ*^2^ *∼ β*. Those sites for which the *CV*^2^ and *F* are both age-dependent, as well as *σ*^2^, were labeled as ‘NA’ group. However, it does not exclude that some other functional dependence *σ*^2^ = *σ*^2^ (*β*) may exist.

The percentage of CpG probes with different normalization types is presented in Figure 7. The number of aDaVMPs with functional dependence *σ*^2^ *∼ β*^2^ (*CV*^2^ *∼ const*) equals 1888 for males and 2021 for females; with functional dependence *σ*^2^ *∼β* (*F∼ const*) it amounts to 129 for males and 238 for females. The number of aDaVMPs for which the *CV*^2^ and *F* retained age-dependence (‘NA’ group) equals 58 for males and 23 for females. The lists of probes for males and females is reported in Supplementary File 11 marked ‘CV2’, ‘Fano’ and ‘NA’. Examples of the most representative CpG sites are given in Figure 8 and Figure 9. The experimental results showed that for most aDaVMPs, the variability increases/decreases as the square of beta value (*CV*^2^ is age independent), that is, *σ*^2^ *∼ β*^2^.

**Figure 7.**
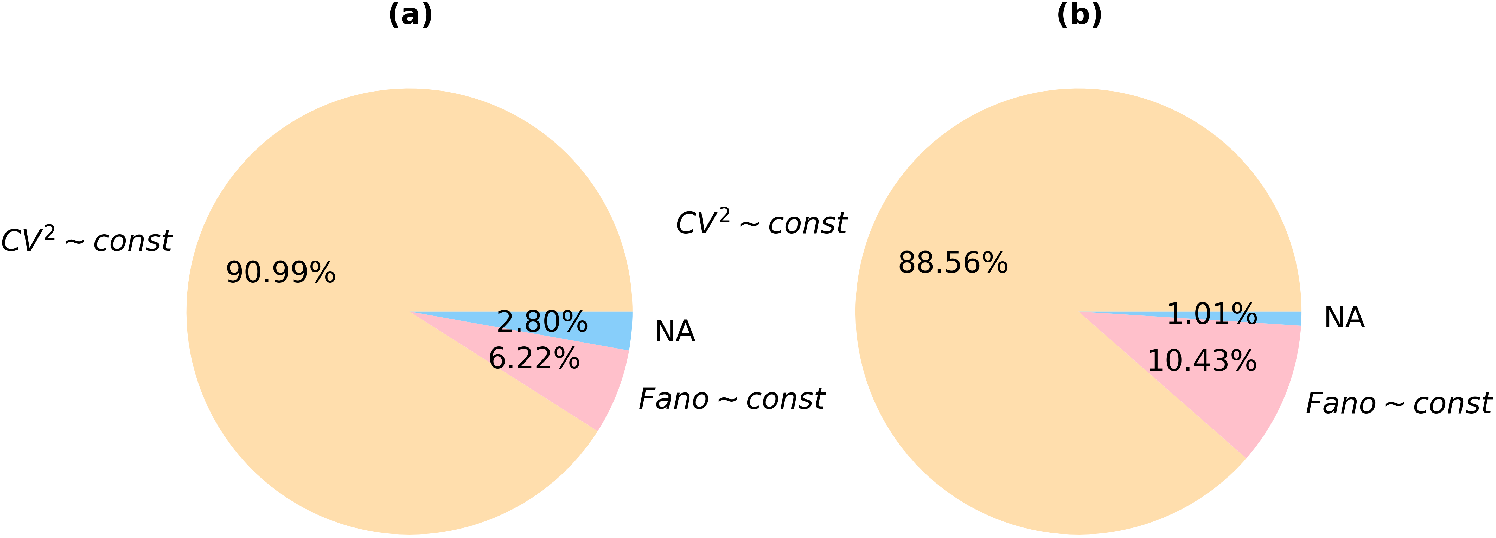
The percentage of aDaVMPs with different normalization types of variability in GSE87571 dataset for males **(a)** and females **(b)**: with functional dependence *σ*^2^ *∼ β*^2^ (*CV*^2^ *∼ const*, beige); with functional dependence *σ*^2^ *∼ β* (*F∼ const*, lightblue); aDaVMPs for which the *CV*^2^ and *F* did not pass by p-value (‘NA’ group, pink).

**Figure 8.**
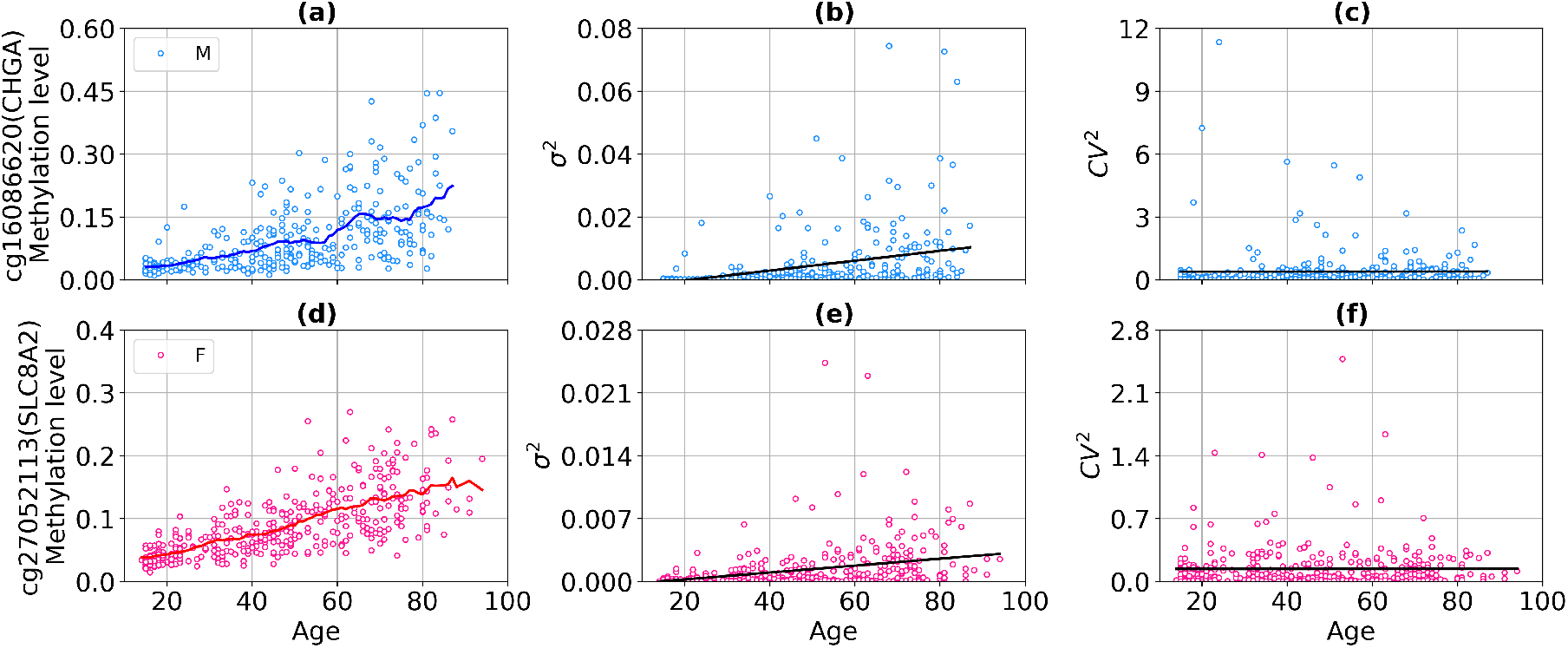
Example of CpG probes for which the squared coefficient of variation (the variability normalized to the squared average beta value) is age independent with a p-value > 0.9 in GSE87571 dataset. The top panel presents scatter plots for males: the methylation level **(a)**, the variability **(b)**, the squared coefficient of variation **(c). (d), (e), (f)** on the bottom panel present similar graphs for females. Curves represent the average level of methylation taken over a sliding window of 10 years for males (blue curves) and females (red curves). Black lines represent linear regression fits.

**Figure 9.**
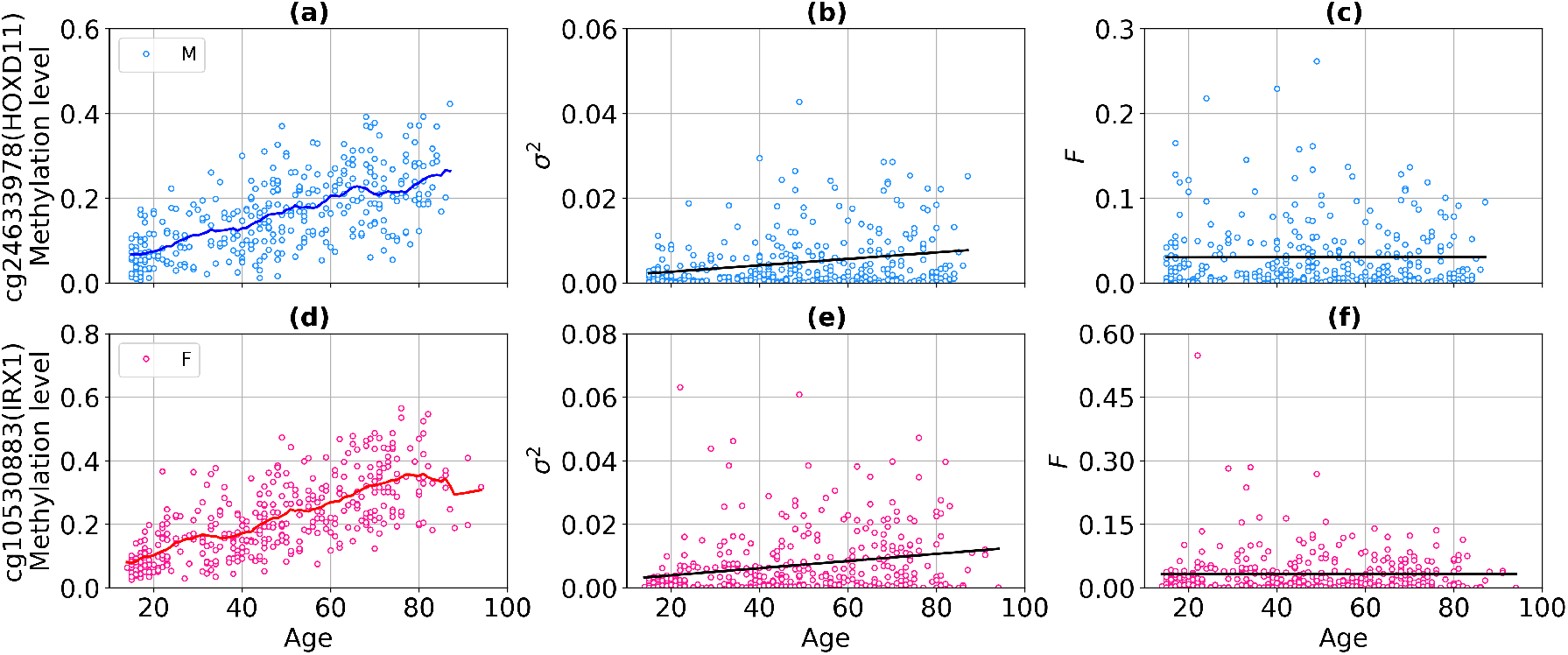
Example of CpG probes for which Fano factor (the variability normalized to the average beta value) is age independent with a p-value > 0.9 in GSE87571 dataset. The top panel presents scatter plots for males: the methylation level **(a)**, the variability **(b)**, the Fano factor **(c). (d), (e), (f)** on the bottom panel present similar graphs for females. Curves represent the average level of methylation taken over a sliding window of 10 years for males (blue curves) and females (red curves). Black lines represent linear regression fits.

Repeating the analysis for the residuals of methylation values, we obtained similar results, in particular, that among 592 and 1008 aDaVMPs sites 89.02% and 86.31% of CpGs have a functional relationship, *σ*^2^ *∼ β*^2^, for males and females, respectively. Probe lists for residuals with division into groups of different functional relationships (*σ*^2^ *∼ β*^2^, *σ*^2^ *∼ β* and ‘NA’) are presented in Supplementary File 12. Common CpGs with functional dependence *σ*^2^ *∼ β*^2^ for beta values and residuals are marked with ‘X’, and common CpGs with functional dependence *σ*^2^ *∼ β* are marked with ‘XX’ in the corresponding Supplementary Files 11, 12.

### Identification of probes having nonlinear age-associated methylation changes

Since in most of CpGs the age-dependent the average methylation and variability are functionally related, it is of particular interest to identify the type of functional dependence. A linear fit, *β ∼ age*, is the most popular model, widely used in epigenetic clocks. Besides, discriminating linear and nonlinear trends requires large age spans. Based on GSE87571 dataset (with age span of 80 years) we aimed to detect probes with average methylation level changing according to the power law, *β ∼ age*^*α*^, or exponentially, *β ∼ e*^*γ·age*^. Note, that both nonlinear models are compatible with the *σ*^2^ *∼ β*^2^ scaling, previously found in the majority of cases.

We consider previously selected 3827 aDMPs for males and 3850 aDMPs for females in GSE87571 dataset. Among them we identified the probes with nonlinear age dependence using the approach described in Methods. The number of found nonlinear CpGs is equal to 255 for males and 97 for females (46 common CpGs). In all cases the quality of power law fit proved to be significantly better than exponential. In less than 8% cases nonlinearity is manifested as saturation towards complete (de)methylation (0 or 1 methylation levels). Typical examples of nonlinear (power law) CpGs are presented in Figure 10. The corresponding lists of power law probes obtained for beta values are presented in Supplementary File 13. GO enrichment analysis of the list of nonlinear probes returned 1 significant biological process in females (leukocyte apoptotic process), which was marginally significant in males.

**Figure 10.**
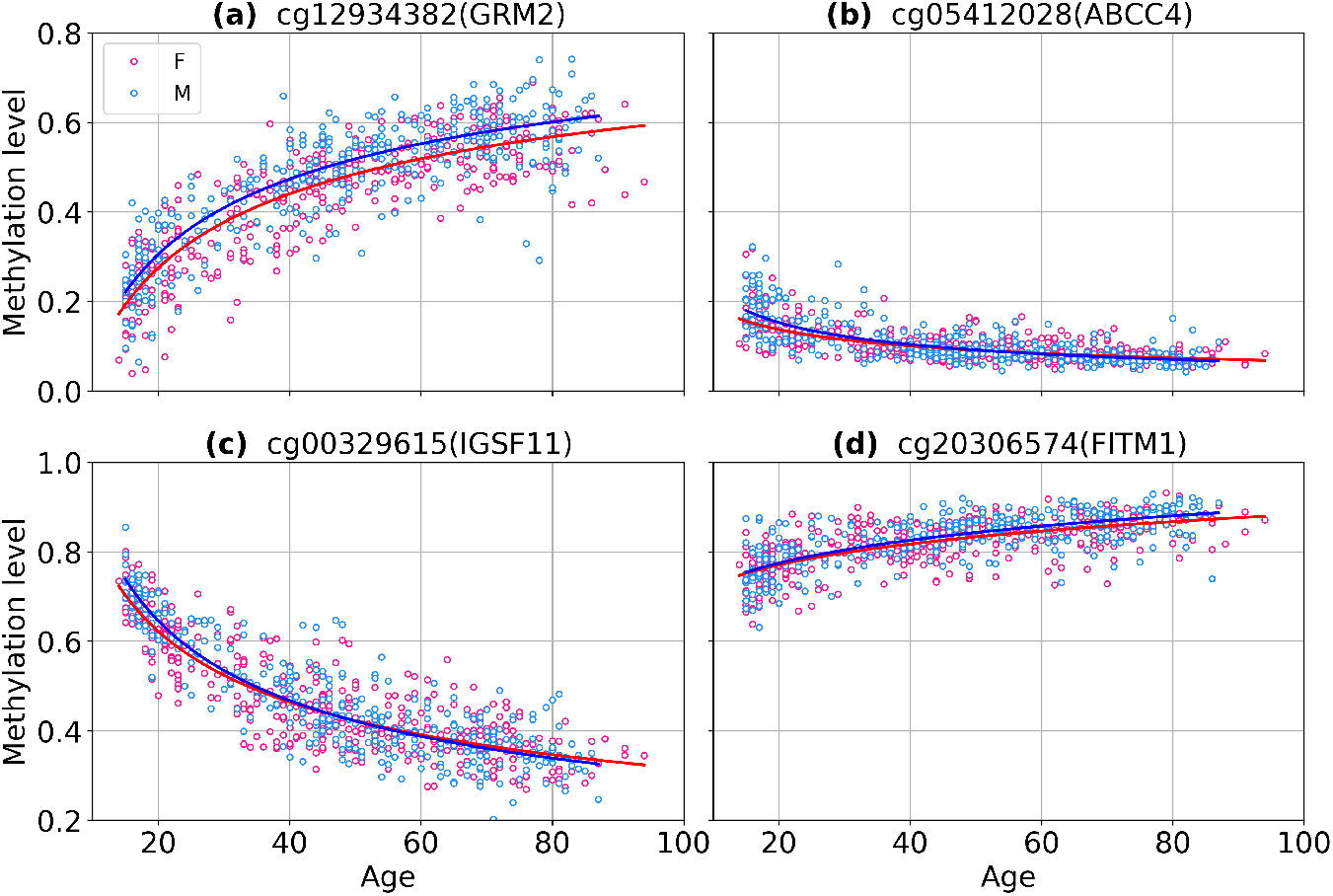
Scatter plots of power law CpGs identified in GSE87571 dataset. Curves represent power law fits *β ∼ age*^*α*^ or 1 *—β ∼ age*^*α*^ for males (blue curves) and females (red curves). **(a), (c)** on the left panel show probes with a methylation level far from the limits, the right panel gives examples of nonlinearity due to saturation to 0 **(b)** and 1 **(d)**.

The number of nonlinear probes for males is considerably greater than for females, that manifests yet another kind of sex specificity of DNA methylation: not simply different in the level of methylation, but having different functional age dependencies, linear and nonlinear. An example of a CpG site with different methylation change laws for males (power law fit) and females (linear fit) is shown in Figure 11.

**Figure 11.**
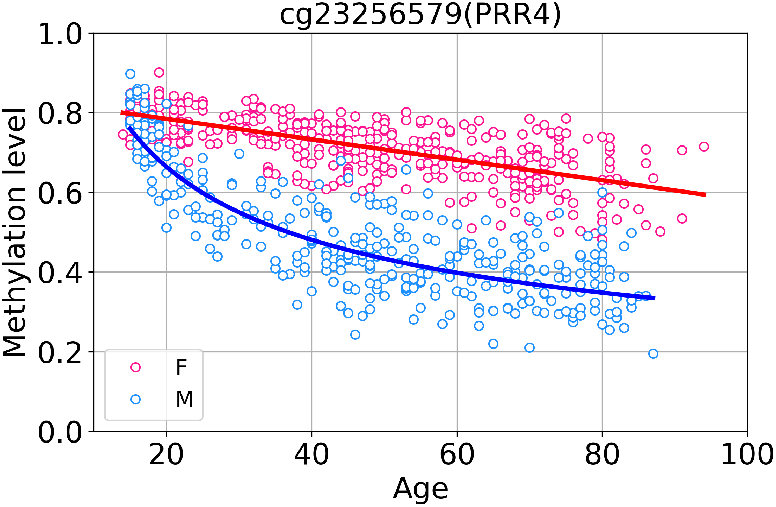
Scatter plot of sex-specific nonlinear CpG cg23256579 (PRR4) in GSE87571 dataset. Curves represent power law fit for males (blue curve) and linear fit for females (red curve).

It follows, that the power law CpGs have a strongly nonlinear dependence of methylation level on age, as the exponents *α* of power fits *β ∼ age*^*α*^ are much less than 1 (cf. the histogram in Figure 12-a). Such nonlinearity dictates that methylation of CpG sites changes faster at young age and slower for the elderly, as compared to an average rate (see Figure 13). Notably, Horvath reported a similar behavior, namely, a nonlinear change in methylation of the 353 Horvath clock CpGs in early childhood^11^. Six CpGs from that set were identified in our analysis too (cg09118625, cg01511567, cg11314684, cg19724470 for males and cg01459453 for females). Importantly, our results demonstrate lifelong nonlinear changes in methylation. Deceleration of DNA methylation change with age was also pointed out^32, 33^.

**Figure 12.**
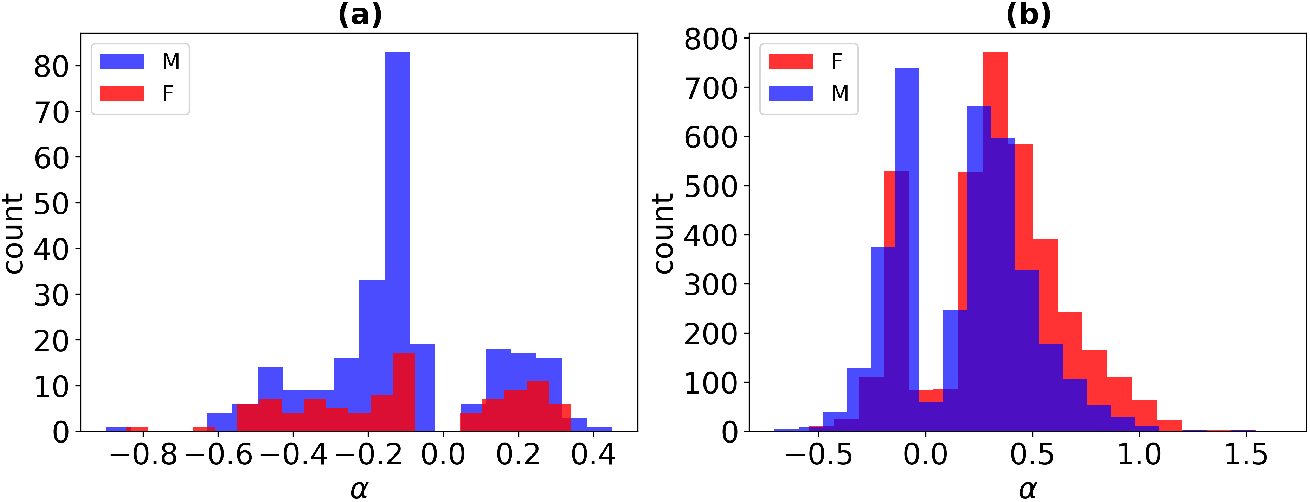
Histogram of the number of males (blue) and females (red) according to exponents *α* of power law fits in GSE87571 dataset: for significantly power law probes **(a)** and for probes for which the power law and linear fits are weakly distinguishable **(b)**.

**Figure 13.**
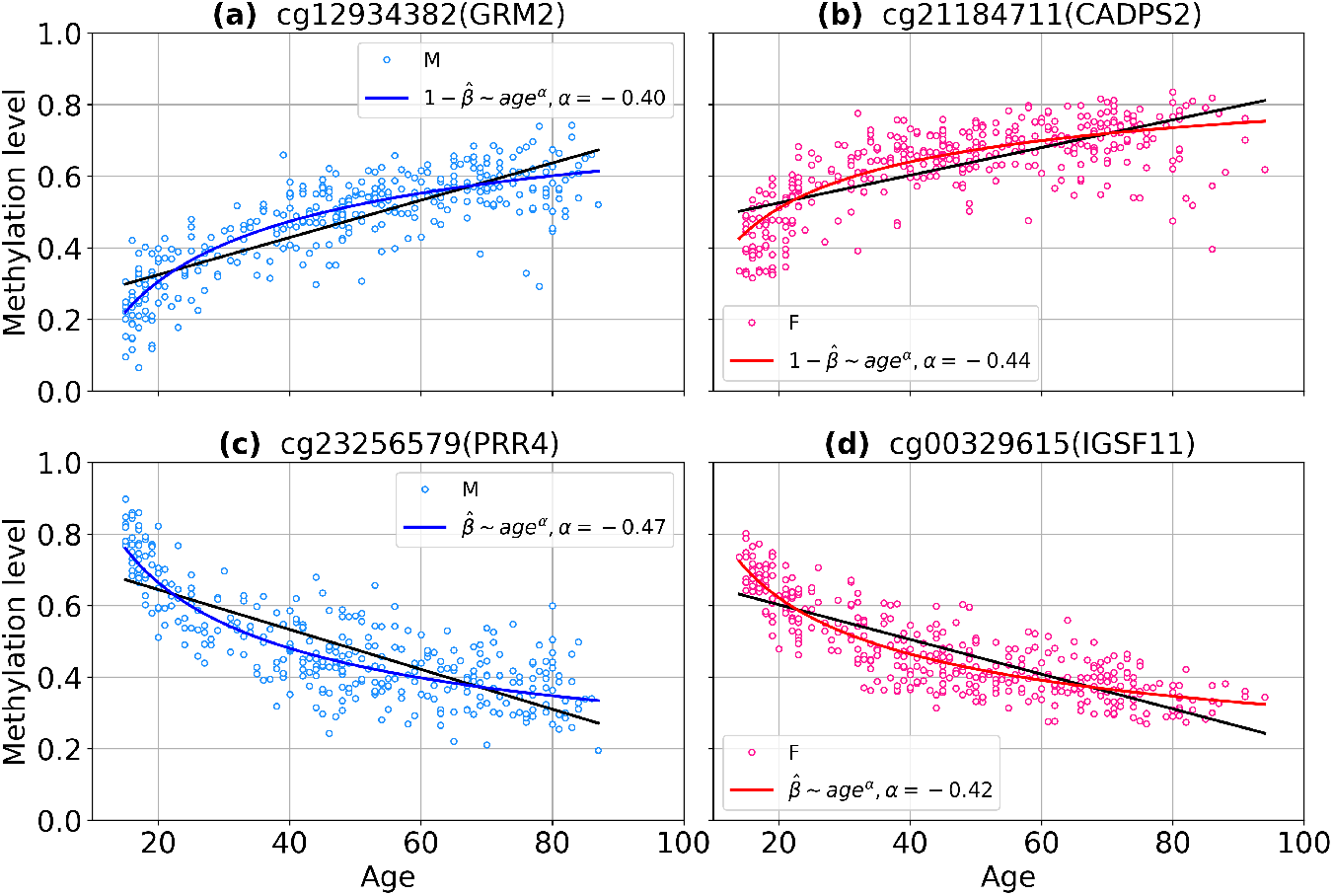
Demonstration of the difference between a power law fits and a linear ones. Scatter plots of power law CpGs identified in GSE87571 dataset. Curves represent a power law fit for males (blue curves) and females (red curves). Black lines represent a linear regression fits. **(a), (c)** on the left panel show methylation level of males, **(b), (d)** on the right panel show methylation level of females.

Interestingly, almost all aDMPs pass the “weak” nonlinearity criterion implying that the power law and linear fits are of comparable quality (cf Materials and Methods for details), amounting to 3564 CpGs for males and 3736 CpGs for females. Moreover, despite the formally close goodness of linear and nonlinear fits, for many of them the visual inspection reveals significant nonlinearity, and the exponents *α* of power fits *β ∼ age*^*α*^ are much less than 1 (see histogram in Figure 12-b). Accordingly, one might expect that the future data, more homogeneous, with even greater age span, and longitudinal would enable more probes to pass the strong nonlinearity criterion.

Nonlinear trends also exist for residuals. Among 2347 aDMPs for males and 2619 aDMPs for females we found 400 and 516 significantly nonlinear CpGs (with power law fit), respectively (Supplementary File 14). CpG sites that are at the intersection of the power law probes lists for beta values and residuals are marked with ‘X’ in the corresponding Supplementary Files 13, 14.

## Discussion

We analyzed deterministic and stochastic sources *D × S* of age-dependent variability of DNA methylation in longitudinal data and demonstrated the dominating role of *D*. For about 90% of the considered CpGs the regular divergence of individual trajectories gives a major contribution to the variability of beta values with age, while the influence of stochastic factors is secondary. Remarkably, the study identified sex differences, as the number of CpG sites with age-dependent variability of methylation and dominating *S* has proved to be greater for males.

The results suggest the leading role of “imprinted” sources of heterogeneity in *G × E* combination that defines individual aging trajectories, such as genetic predisposition, early development, and robust environmental factors (socio-economical status, lifestyle and habits, climate and ecological niches), while random perturbations such as genetic noise and short-term environmental changes seem to play a minor role.

Our results are consistent with the recent findings on genetic and environmental contributions to DNA methylation^8^. Reynolds at al. examined two cohorts of longitudinal data: Swedish and Danish twins across the 10-year span (about 68-78 years). The analysis was based on building bivariate biometrical twin models for M-values, for which the relative similarity of monozygotic versus dizygotic twins was compared. Reynolds et al. found that the age-related CpG sites, as well as the sites constituting Hannum, Horvath, Levine and Zhang epigenetic clocks showed a significantly higher proportion of variability attributed to heritable and shared environmental influences due to stronger genetic and common environmental influences. In addition, the authors identified two groups of CpG sites: the so-called “high heritability” and “low stability”. The methylation of the 5037 high heritability CpGs would be stable in time, and manifest variability due to genetic and common environmental influences. The 2020 low stability sites meanwhile showed increased variability across time due to non-shared environmental factors. Our study that made use of the mixed longitudinal and cross-sectional setup and analyzed the signal-to-noise ratio gave corroborating results. The sites that we identify as deterministic can showcase increased methylation variability in time due to genetic and common environmental factors, analogously to high-heritability CpGs. The sites identified as stochastic, where the variability increases due to non-shared environmental influences and genetic noise, may be relate to the low stability CpGs. It has to be noted, that there is no direct identity of these classes, since, although we considered the change in variability between twins, which obviously have a common genetic background and common environmental conditions at least in childhood, our approach does not discriminate genetic and environmental influences. This limitation, however, is recouped by the increased timespan and the number of longitudinal datapoints for a single individual, enabling the signal-to-noise calculation.

The detailed study of the methylation variability confirmed the previously shown predominant increase in variability with age. It was found that there is a mathematical functional relationship between the time-dependence of the methylation level and variability. For about 90% aDaVMPs, the variability changes nonlinearly as *σ*^2^ *∼ β*^2^, whereas for about 6-10%, depending on sex, it follows the linear dependence *σ*^2^ *∼ β*, and only for the few percent cases the dependence remained undetermined.

The regression analysis has revealed significantly nonlinear (power law) epigenetic biomarkers of aging. According to the results, males have more nonlinear probes than females. This adds a novel evidence of the sex specificity of DNA methylation. A significant number of probes demonstrate a nonlinear change in methylation during aging with accelerating for young and slowing down for the elderly people. These nonlinear aDMPs can serve as the basis for the construction of new epigenetic clocks that take into account nonlinear changes in methylation.

## Acknowledgements

Authors acknowledge support by the grant of the Ministry of Education and Science of the Russian Federation Agreement No. 074-02-2018-330. A.Z. acknowledge support by the MRC grant MR/R02524X/1.

## Author contributions statement

C.F., M.I., A.Z. designed the study, M.I., O.V., M.G.B. developed methodology of detecting deterministic / stochastic / nonlinear probes and analysed the results, O.V. conducted the experiments, M.G.B. performed the GO enrichment analysis. All authors have written and reviewed the manuscript.

## Additional information

Authors declare no competing interests.

