## Supplementary Materials for "Disentangling age-dependent DNA methylation: deterministic, stochastic, and nonlinear"

### Supplementary Figures

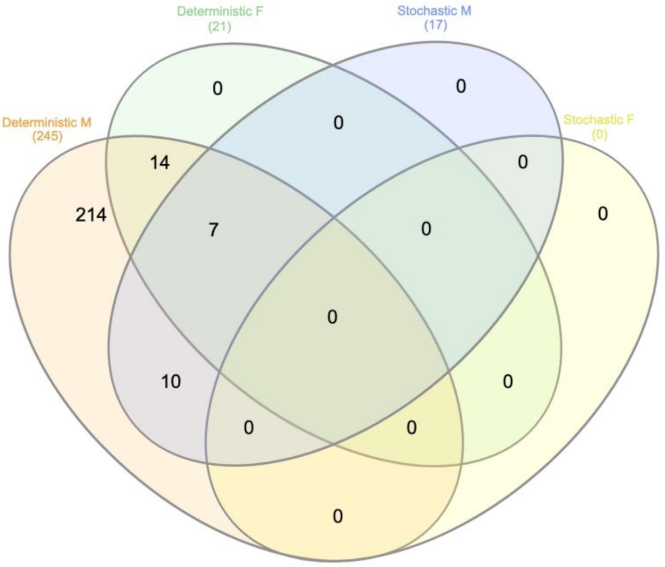


**Supplementary Figure 1.** Venn Diagram of the significantly enriched GO in deterministic and stochastic aVMPs identified in males (M) and females (F).

### Supplementary Files

**Supplementary File 1.** Lists of age-associated variably methylated positions, aVMPs, (column *"name_cpg"*) computed for beta values. These lists were compared with similar lists obtained for residuals of methylation values (Supplementary File 3, intersecting probes are marked with "X" in the *"isIntersect"* column).

**Supplementary File 2.** Gene Ontology enrichment of aVMPs lists (obtained for beta values).

**Supplementary File 3.** Lists of age-associated variably methylated positions, aVMPs, (column *"name_cpg"*) computed for residuals of methylation values. These lists were compared with similar lists obtained for beta values (Supplementary File 1, intersecting probes are marked with "X" in the *"isIntersect"* column).

**Supplementary File 4.** Lists of age-associated variably methylated positions, aVMPs, computed for beta values (column *"name_cpg"*) divided into deterministic and stochastic probes (column *"type"*). Subsets of deterministic and stochastic CpGs were compared with similar subsets obtained for residuals of methylation values (Supplementary File 6, common deterministic CpGs for beta values and residuals are marked with "X" and common stochastic CpGs are marked with "XX" in the column *"isIntersect"*).

**Supplementary File 5.** Gene Ontology enrichment of deterministic and stochastic lists of probes (obtained for beta values).

**Supplementary File 6.** Lists of age-associated variably methylated positions, aVMPs, computed for residuals of methylation values (column *"name_cpg"*) divided into deterministic and stochastic probes (column *"type"*). Subsets of deterministic and stochastic CpGs were compared with similar subsets obtained for beta values (Supplementary File 4, common deterministic CpGs for beta values and residuals are marked with "X" and common stochastic CpGs are marked with "XX" in the column *"isIntersect"*).

**Supplementary File 7.** Lists of age-associated differentially methylated positions, aDMPs, (column *"name_cpg"*) computed for beta values. These lists were compared with similar lists obtained for residuals of methylation values (Supplementary File 9, intersecting probes are marked with "X" in the column *"isIntersect"*).

**Supplementary File 8.** Lists of age-associated differentially and variably methylated positions, aDaVMPs, (column *"name_cpg"*) computed for beta values. These lists were compared with similar lists obtained for residuals of methylation values (Supplementary File 10, intersecting probes are marked with "X" in the column *"isIntersect"*).

**Supplementary File 9.** Lists of age-associated differentially methylated positions, aDMPs, (column *"name_cpg"*) computed for residuals of methylation values. These lists were compared with similar lists obtained for beta values (Supplementary File 7, intersecting probes are marked with "X" in the column *"isIntersect"*).

**Supplementary File 10.** Lists of age-associated differentially and variably methylated positions, aDaVMPs, (column *"name_cpg"*) computed for residuals of methylation values. These lists were compared with similar lists obtained for beta values (Supplementary File 8, intersecting probes are marked with "X" in the column *"isIntersect"*).

**Supplementary File 11.** Lists of age-associated differentially and variably methylated positions, aDaVMPs, computed for beta values (column *"name_cpg"*) divided into three functional groups: "CV2" ($\sigma^2 \sim \beta^2$), "Fano" ($\sigma^2 \sim \beta$) and "NA" (in column *"type"*). Subsets of "CV2" and "Fano" CpGs were compared with a similar subsets obtained for residuals of methylation values (Supplementary File 12, common "CV2" CpGs for beta values and residuals are marked with "X" and common "Fano" CpGs are marked with "XX" in the column *"isIntersect"*).

**Supplementary File 12.** Lists of age-associated differentially and variably methylated positions, aDaVMPs, computed for residuals of methylation values (column *"name_cpg"*) divided into three functional groups: "CV2" ($\sigma^2 \sim \beta^2$), "Fano" ($\sigma^2 \sim \beta$) and "NA" (in column *"type"*). Subsets of "CV2" and "Fano" CpGs were compared with a similar subsets obtained for beta values (Supplementary File 11, common "CV2" CpGs for beta values and residuals are marked with "X" and common "Fano" CpGs are marked with "XX" in the column *"isIntersect"*).

**Supplementary File 13.** Lists of CpGs with nonlinear (power law) methylation change (column *"name_cpg"*) computed for beta values. These lists were compared with similar lists obtained for residuals of methylation values (Supplementary File 14, intersecting probes are marked with "X" in the column *"isIntersect"*).

**Supplementary File 14.** Lists of CpGs with nonlinear (power law) methylation change (column *"name_cpg"*) computed for residuals of methylation values. These lists were compared with similar lists obtained for beta values (Supplementary File 13, intersecting probes are marked with "X" in the column *"isIntersect"*).
